# The loss of enzymatic activity of the PHARC associated lipase ABHD12 results in increased phagocytosis that causes neuroinflammation

**DOI:** 10.1101/2021.08.22.457059

**Authors:** Shubham Singh, Siddhesh S. Kamat

**Affiliations:** Department of Biology, Indian Institute of Science Education and Research (IISER) Pune, Dr. Homi Bhabha Road, Pashan, Pune 411008, Maharashtra, India

## Abstract

Phagocytosis is an important evolutionary conserved process, essential for clearing pathogens and cellular debris in higher organisms, including humans. This well-orchestrated innate immunological response is intricately regulated by numerous cellular factors, important amongst which, are the immunomodulatory lysophosphatidylserines (lyso-PSs) and the pro-apoptotic oxidized phosphatidylserines (PSs) signaling lipids. Interestingly, in mammals, both these signaling lipids are physiologically regulated by the lipase ABHD12, mutations of which, cause the human neurological disorder PHARC. Despite the biomedical significance of this lipase, detailed mechanistic studies and the specific contribution of ABHD12 to innate processes like phagocytosis remain poorly understood. Here, by immunohistochemical and immunofluorescence approaches, using the murine model of PHARC, we show, that upon an inflammatory stimulus, activated microglial cells in the cerebellum of mice deficient in ABHD12 have an amoeboid morphology, increased soma size, and display heightened phagocytosis activity. We also report that upon an inflammatory stimulus, cerebellar levels of ABHD12 increase to possibly metabolize the heightened oxidized PS levels, temper phagocytosis and in turn control neuroinflammation during oxidative stress. Next, to complement these findings, using biochemical approaches in cultured microglial cells, we show that the pharmacological inhibition and/or genetic deletion of ABHD12 results in increased phagocytic uptake in a fluorescent bead uptake assay. Together, our studies provide compelling evidence that ABHD12 plays an important role in regulating phagocytosis in cerebellar microglial cells, and provides a possible explanation, as to why human PHARC subjects display neuroinflammation and atrophy in the cerebellum.

## INTRODUCTION

Phagocytosis is an important innate immune response, conserved in both prokaryotes and eukaryotes^*1*^. In early single cellular life forms, phagocytosis is an important way to obtain nutrients, while, in more evolved complex multicellular organisms, this physiologically indispensable response, is central to an organism’s immunity^*1, 2*^. In higher organisms, phagocytosis is the first type of defense against invading pathogens, and is essential for clearing infections, or cellular debris^*1, 3, 4*^. In mammals (including humans), the phagocytic cells (typically macrophages or cells with a monocyte lineage) have an innate ability to recognize, engulf and digest foreign solid particles ≥ 500 nm (e.g. pathogens like viruses and bacteria) or any defective/toxic material generated during heightened metabolic states and/or injury (e.g. diseased cells, protein aggregates, tissue debris) with sufficient antigenicity^*3-5*^. Often, a distressed (injured or infected or apoptotic) cell needs to be cleared from the circulatory system, to prevent any damage to its surrounding cells or tissues, and such a cell secretes substances (e.g. signaling lipids, chemokines) that provide phagocytic cells (e.g. circulating macrophages) signature signaling cues, to initiate its engulfment and clearance from the system via phagocytosis^*2, 3, 6-8*^. Amongst the signaling lipids thought to be involved in facilitating phagocytosis^*1, 2, 4*^, are the immunomodulatory lysophosphatidylserines (lyso-PSs) ^*9*^ and the pro-apoptotic oxidized phosphatidylserines (PSs)^*10*^.

Of the many important immunological functions that the lyso-PSs perform in mammalian physiology^*9*^, this signaling lipid in particular, promotes efferocytosis^*6, 11, 12*^, and plays an important role in the resolution of inflammation by enhancing the phagocytic activity of macrophages^*9, 13, 14*^. In mammalian macrophages, we have recently shown that the very long chain (VLC) lyso-PSs activate these innate immune cells and elicit the secretion of pro-inflammatory cytokines (e.g. tumor necrosis factor alpha (TNF-α) and interleukin 6 (IL-6)) via a Toll-like receptor 2 (TLR2) dependent pathway, while long chain (LC) lyso-PSs regulate various signaling cascades (cytosolic calcium influx, cyclic adenosine monophosphate production and phosphorylation of the nodal signaling protein extracellular signal-regulated kinase (ERK)) possibly through an as-of-yet unknown G-Protein Coupled Receptor (GPCR)^*15*^. On the other hand, the pro-apoptotic oxidized PSs have a long-standing association with phagocytosis, particularly in the context of clearing up diseased/dying cells during oxidative stress or tissue injury^*16-19*^. Given the asymmetric localization of almost all the PS to the inner leaflet of the membrane bilayer, the externalization of PS (due to its non-enzymatic oxidation to yield oxidized PSs), represents a cell in oxidative stress and undergoing apoptosis^*20, 21*^. This extracellularly facing PS from such distressed cells is an “eat me” cue to circulating macrophages, whose scavenger receptors (e.g. CD36) recognize this externalized oxidized PS, and initiate signaling pathways to start engulfing and digesting these apoptotic cells via phagocytosis^*20, 21*^.

Interestingly, in all mammals (including humans), the integral membrane lipase ABHD12 (α/β-hydrolase domain containing protein # 12) from the metabolic serine hydrolase family^*22*^, controls the physiological concentrations of both lyso-PSs^*23, 24*^ and oxidized PSs^*10*^ in the central nervous system and various immune cells, including macrophages, particularly during conditions of oxidative stress (**Figure 1**). This metabolic activity of ABHD12 has significant biomedical importance, as deleterious (null) mutations to the gene encoding this lipase in humans, results in an autosomal recessive early onset neurodegenerative disease called PHARC (***<u>p</u>***olyneuropathy, ***<u>h</u>***earing loss, ***<u>a</u>***taxia, ***<u>r</u>***etinitis pigmentosa and ***<u>c</u>***ataract) (**Figure 1**)^*25, 26*^. Of note, immunohistochemical and behavioral studies performed in ABHD12 knockout mice, the murine model of PHARC, show that the loss of enzymatic activity of ABHD12 is associated with an increase in the number of activated microglia (brain-resident macrophage) and in turn neuroinflammation in the cerebellum (in the arbor vitae region), that eventually manifests in neurobehavioral defects (e.g. auditory and motor deficiencies) closely mimicking those observed in human PHARC subjects^*24*^. Additionally, lipidomics studies from cultured mammalian cells following the pharmacological and/or genetic disruption of ABHD12 have shown, that the loss of ABHD12 activity causes the accumulation of both lyso-PSs and oxidized PSs, that results in increased activation of macrophages devoid of ABHD12 relative to wild-type controls^*10, 23*^.

**Figure 1.**
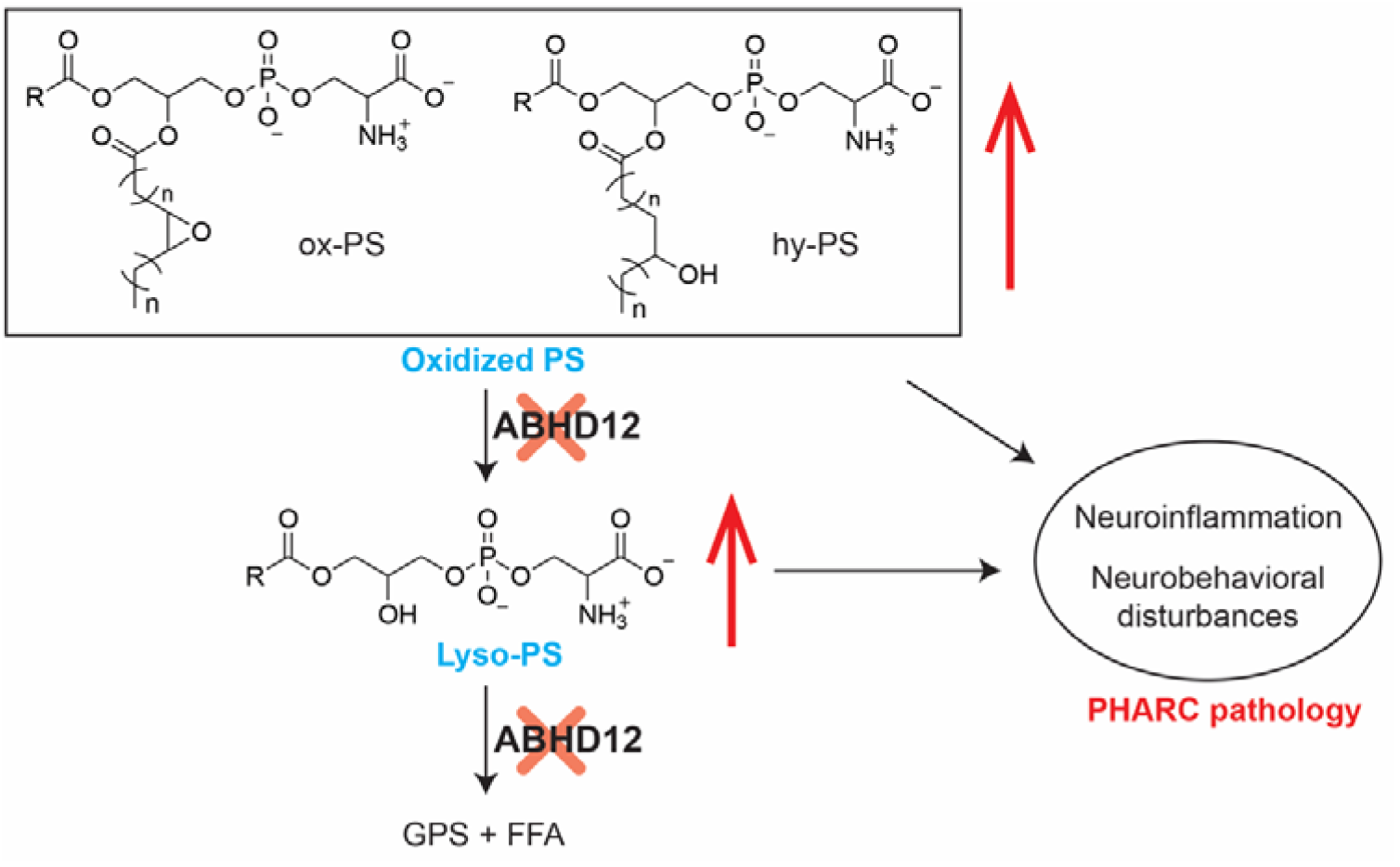
The biochemical pathways regulated by the lipase ABHD12. Loss of ABHD12 lipase activity causes accumulation of lyso-PS^*24*^ and oxidized PS lipids^*10*^ in the brain, that contribute to neuroinflammation, that manifests into the neurobehavioral defects associated with the human neurological disorder PHARC^*25*^. GPS = glycerophosphoserine and FFA = free fatty acid. This figure is adapted with modifications from a previous study from our lab^*10*^.

Given that the enzymatic activity of ABHD12 controls physiological concentrations of the signaling lipids that are associated with the process of phagocytosis^*9*^, we wanted to assess if the loss of this lipase activity has any effect on phagocytosis, and if this has any implications on our understanding of the pathology of the human neurological disorder PHARC^*25*^. Towards this, using immunohistochemical, immunofluorescence and biochemical approaches in ABHD12-null mice and microglial cell lines, we show that the loss of ABHD12 activity results in heightened phagocytosis activity by (activated) microglial cells, and our studies taken together, perhaps provides a preliminary explanation, as to why aged ABHD12 knockout mice display neuroinflammation and human PHARC subjects show atrophy in their cerebellum.

## MATERIALS AND METHODS

### Materials

Unless mentioned otherwise, all chemicals, buffers and reagents described in this paper were purchased from Sigma-Aldrich. All mammalian tissue culture media, consumables, antibiotics and fetal bovine serum (FBS) were purchased from HiMedia. All primary and secondary antibodies were purchased from Abcam and Thermo Fisher Scientific respectively, unless stated otherwise. The fluorescent carboxylate modified microspheres (fluorescent latex beads) (Thermo Fisher Scientific, Catalog # F8813) were a kind gift from Prof. Roop Mallik (IIT Mumbai). The ABHD12 selective inhibitor JJH350 used in this study was a kind gift from Prof. Benjamin F. Cravatt (Scripps Research).

### Animal studies

All animal (mouse) studies described in this paper have received formal approval from Institutional Animal Ethics Committee at IISER Pune (IAEC–IISER Pune) (application numbers: IISER_Pune IAEC/2019_02/07 and IISER_Pune IAEC/2021_01/09), constituted as per the guidelines outlined by the Committee for the Purpose of Control and Supervision of Experiments on Animals (CPCSEA), Government of India. All mice were bred and maintained at National Facility for Gene Function in Health and Disease (NFGFHD), IISER Pune. The ABHD12 knockout mice used in this study were generated on the C57BL/6-genetic background as reported earlier^*24*^. All mice used in the study were generated by a standard ABHD12 heterozygous x ABHD12 heterozygous breeding scheme, genotyped using previously established protocols^*24*^, and had *ad libitum* access to food and water. For all the animal studies described here, an equal number of age-matched male and female mice (10 – 12 weeks of age) were used to control for any gender and age specific effects. Prof. Benjamin F. Cravatt (Scripps Research) is thanked for generously gifting the ABHD12 heterozygous breeder mice used to generate the ABHD12 knockout mice used in this study.

### Immunohistochemical and immunofluorescence sample preparations

All immunohistochemical (IHC) experiments and analysis were done using protocols recently reported by us^*15, 27*^. Briefly, wild type or ABHD12 knockout mice (10−12 weeks old, equal number of males and females used in any given experiment) were deeply anaesthetized using isoflurane and perfused, first with chilled phosphate-buffered saline (PBS) and then with ice cold 4% (w/v) paraformaldehyde (PFA) in PBS. To ensure anatomical integrity, the brains were gently harvested, and transferred to 4% (w/v) PFA in PBS overnight (∼ 16 – 18 h) at 4 °C to ensure proper tissue fixation. Thereafter, the fixed brains were transferred into a 30% (w/v) sucrose solution in PBS and maintained at 4 °C until the brains sank to the bottom of the tubes (∼ 3 days). Cryoprotected brains were sectioned in 25 μm thick coronal sections on a Leica CM1950 cryotome maintained at −30 °C. Every 5^th^ cerebellar section was collected and stored in PBS at 4 °C, followed by its processing for IHC experiments. For DAB-based bright-field IHC analysis, endogenous peroxidases were inactivated and/or quenched with 3% (v/v) hydrogen peroxide in PBS for 15 min at 25 °C in the dark, following which, the sections were washed twice with 1% (w/v) bovine serum albumin (BSA) in PBS and permeabilized with 0.1% (w/v) Triton X-100 and 0.5% (w/v) BSA in PBS for 45 min at 25 °C. Following this, the sections were incubated with the primary antibody (dilution 1:100) in 1% (w/v) BSA overnight (∼ 12 – 16 hours) at 4 °C, washed three times with 0.5% (w/v) BSA in PBS and incubated with the secondary antibody, biotinylated horse anti-rabbit (Vector laboratories, BP-1100) or biotinylated goat anti-mouse (Vector laboratories, BP-9200) at a dilution of 1:1000, in 0.5% (w/v) BSA in PBS for 1 hour at 25 °C. Following this, the sections were washed thrice in 0.5% (w/v) BSA in PBS, incubated with ABC Elite Vectastatin (Vector laboratories, PK-6100) for 1 hour at 25 °C, and washed thrice with excess PBS at 25 °C. Finally, the sections were stained with ImmPACT DAB (Vector laboratories, SK-4105) in the dark for 3 mins, and washed five times with PBS to remove excess DAB. Thereafter, the slides were mounted using VectaMount (Vector laboratories, H-5000). All the stained sections were imaged using under bright field channel using a Zeiss Apotome microscope. For fluorescence IHC analysis, comparable cerebellar sections were permeabilized with 0.1% (v/v) Triton X-100 and 0.5% (w/v) BSA in PBS for 45 mins at 25 °C and blocked with 1% (w/v) BSA in PBS for 60 mins at 25 °C. Following this step, the sections were washed thrice with 0.5% (w/v) BSA in PBS and incubated overnight (∼ 12 – 16 hours) with the primary antibody (dilution 1:100) at 25 °C. Thereafter, the sections were washed thrice with 0.5% (w/v) BSA in PBS and incubated with the secondary antibody (1:1000 dilution) for 60 mins at 25 °C. Sections were again washed with PBS (three times) and mounted with Fluoromount aqueous mounting medium (Sigma-Aldrich, F4680). The stained sections were imaged using a Leica SP8 confocal laser scanning microscope. The primary antibodies used in these studies were: anti-ABHD12 (rabbit, monoclonal, Abcam, catalog ab182011); anti-Iba1 (rabbit, monoclonal, Abcam, catalog ab178846), anti-Iba1 (mouse, monoclonal, Santa Cruz Biotechnology, catalog sc-32725), and anti-LAMP1 (rabbit, polyclonal, Abcam, catalog ab24170). The following secondary antibodies were used in this study were: goat anti-rabbit IgG (H&L) conjugated with DyLight 488 (Thermo Fisher Scientific, 35553) and goat anti-mouse IgG (H&L) conjugated with Alexa Fluor 568 (Thermo Fisher Scientific, A-11004).

### Activated microglia counting and analysis in the IHC experiments

Using the Fiji-ImageJ1.50i [National Institutes of Health (NIH), for Windows OS] software, all the 2-D bright field images were converted to 8-bit grey scale (Image<<Type<<8-bit). The thresholding for these grey scale images were manually done to mark the soma of the microglia (Image<<Adjust<<Threshold), and the soma size (area) of the microglia were measured using particle analysis tool (Analyze>>Analyze Particles) with set output of area of the particles and their number. From the Iba-1 immunostaining profiles, the microglia with soma size (area) > 200 μm^2^ were considered as activated as per previous reports^*15, 24*^, while those with soma size < 200 μm^2^ were considered non-activated. For size comparisons between the two genotypes, soma size of all the activated microglia in both wild type and ABHD12 knockout mice from the different biological replicates were used in the analysis. Based on previous results for the neuroinflammation observed in ABHD12 null mice, the arbor vitae region near deep cerebellar nuclei in the cerebellum were used for this IHC analysis, because this region also has cleaner microglia staining with very minimal background of DAB, and the microglial morphology and number of microglial cells is also largely homogenous with less variation basally, compared to other anatomical regions in cerebellum (e.g. the granular layers or molecular layers of different cerebellar peduncles).

### Quantification of fluorescent markers in the IHC experiments

For all the IHC experiments, 3-D stacks were imaged and recorded on a confocal microscope and all analysis was performed offline using Fiji-ImageJ1.50i [National Institutes of Health (NIH), for Windows OS] software. For quantitation, the image stack showing middle plane of the cells with visible nuclei in center and least overlap of the microglia cytoplasmic marker (Iba1) with nucleus was chosen manually and saved as a colored (RGB) image. Cell boundaries of microglia were marked manually using the Iba1 signal as reference, following which, the channel of interest was extracted (green for LAMP1 or ABHD12), and the images of extracted channel were converted to 8-bit scale. Thereafter, intensity of the protein of interest was quantified for the whole cell by manually selecting the cell boundary. In this experiment, at least 10 such cells were counted per mice (i.e. per biological replicates) and the average (normalized) intensity per mouse (i.e. biological replicate) was reported. As described previously, this analysis was performed in the arbor vitae region of the cerebellum, which also has the least fluorescent noise and the microglia of this anatomical region are morphologically homogenous.

### Western Blot Analysis

All western blot analysis (immunoblotting) was done using protocols previously reported by us^*27, 28*^. Briefly, the protein lysates (50 μg) were resolved on a 12.5% SDS−PAGE gel, and transferred onto a methanol-activated PVDF membrane (GE Healthcare) at 60 V for 12 h at 4 °C. Following the transfer, the membrane was incubated with the respective primary and secondary antibodies at 1:1000 and 1:10000 dilutions, respectively using standard protocols reported by us^*27, 28*^. The blots were developed using the Thermo West Pico Western blotting substrate (Thermo Fisher Scientific) and the images were recorded on a Syngene G-Box Chemi-XRQ gel documentation system. The following primary antibodies were used in this study: anti-ABHD16A (rabbit, monoclonal, Abcam, 185549), anti-ABHD12 (rabbit, monoclonal, Abcam, ab182011), anti-Iba1 (mouse, monoclonal, Santa Cruz Biotechnology, catalog sc-32725), anti-GAPDH (rabbit, monoclonal, Abcam, catalog ab1818602), and anti-β-actin (rabbit, polyclonal, Cloud Clone Corp., CAB340Hu01). The following secondary antibodies were used in this study: HRP-conjugated anti-rabbit IgG (goat, Thermo Fisher Scientific, 31460) and HRP-linked anti-mouse IgG (goat, Cloud Clone Corp., SAA544Mu19). All densitometric analyses of bands detected in these immunoblotting experiments were performed using Fiji-ImageJ 1.50i [National Institutes of Health (NIH), for Windows OS] software^*29, 30*^. To quantify the blots, lanes on the acquired western blot images were manually marked and selected, and intensity of the band were determined using “Plot Lanes” option available under Analyse>>Gels in the software. Thereafter, the area under the curve (for intensity plot) was measured using the wand tracing tools. All intensities of proteins of interest were divided by intensity of marker control proteins per experiment, per replicate, and the ratios obtained from this, were graphically plotted.

### Mammalian cell lines culturing

The BV-2 (RRID: CVCL_0182) and N9 (RRID: CVCL_0452) immortalized microglial cell lines were a kind gift from Prof. Anirban Basu (National Brain Research Centre, India). Both the cell lines were stained routinely with DAPI to ensure that they were devoid of any mycoplasma contamination. The N9 microglial cell line was cultured in RPMI1640 media supplemented with 10% (v/v) heat inactivated FBS, and 1X antibiotics (penicillin-streptomycin, MP Biomedicals) at 37 °C with 5% (v/v) CO_2_. The BV-2 microglial cell line was cultured in DMEM media supplemented with 10% (v/v) heat inactivated FBS, and 1X antibiotics (penicillin-streptomycin, MP Biomedicals) at 37 °C with 5% (v/v) CO_2_. Primary microglia cells were isolated from brains of day 18 (E18) embryos of mice as reported previously. Briefly, brains were dissected out of E18 embryos, meninges were carefully stripped and removed. Thereafter, the brain was trypsinized using 0.25% (w/v) trypsin on ice for 10 mins, followed by incubation with DNAse I (HiMedia - ML068) for another 5 mins on ice. Following this, the cells of the brain were pelleted at 300 g for 15 mins at 4 °C, suspended in DMEM media by pipetting and plated on poly-D-lysine coated glass coverslips. The cells were maintained for 3-5 days in DMEM media supplemented with 10% (v/v) heat inactivated FBS, and 1X antibiotic-antimycotic (Thermo Fisher Scientific, catalog # 15240096) at 37 °C with 5% (v/v) CO_2_. The media for this microglial cell culture was changed every 2 days and microglia cells were identified by anti-Iba1 immunostaining. Primary thioglycollate elicited peritoneal macrophages were generated from mice, and cultured using established protocols recently reported by us^*10, 27*^. For all the cellular experiments, the cells were counted using a Trypan Blue staining method on a BC20 automated cell counter (Bio-Rad) as per manufacturer’s instructions.

### Phagocytosis assay

To measure the extent of phagocytosis by microglial cells and/or macrophages, established fluorescent latex bead engulfment assays were performed. For this bead engulfment assay, microglial cells (BV-2 or N9) or primary microglia or macrophages were grown on poly-D-lysine coated glass coverslips to about 50 – 60% confluency. Thereafter, these cells were incubated with fluorescent carboxylate modified microspheres (fluorescent latex beads, 0.5 μm, Thermo Fisher Scientific, Catalog # F8813) at a 1:200 cell to bead ratio, for 30 minutes at 37°C, 5% (v/v) CO_2_. Following the bead incubation step, the cells were washed with ice-cold sterile Dulbecco’s Phosphate Buffered Saline (DPBS) and fixed with 4% (w/v) paraformaldehyde (PFA) in DBPS at ambient temperature (23 – 25 °C) using established protocols^*27, 28*^, and subsequently imaged to count the number of beads engulfed by phagocytosis by the microglial cells and/or macrophages. The *N*-hydroxyhydantoin-carbamate JJH350, a ABHD12 specific covalent inhibitor^*31*^, was used for the pharmacological inhibition of ABHD12 in the bead engulfment assays, where the cells were treated with 10 μM JJH350 for 6 hours, following which the bead engulfment assay was performed as described above. All quantifications and image analysis were done offline using Fiji-ImageJ1.50i [National Institutes of Health (NIH), for Windows OS] software. For quantitation, image stack showing middle plane of the cell with visible nuclei in center and least overlap of the cellular cytoplasmic marker (Iba1 and/or Phalloidin) with nucleus was chosen manually and saved as a colored RGB image. Cell boundaries of microglia were marked manually using the Phalloidin signal as a reference, following which, the channel of interest was extracted (green for beads), and the resulting images of extracted channel were converted to 8-bit scale. Thereafter, the intensity threshold was manually adjusted using the software route (Image>>Color>>Threshold) to mark the beads, and these engulfed particles were analyzed using analyze particles option (Analyze>>Analyze Particles) with print option of particle size (area) and number. This analysis gave us the number of beads that were engulfed per cell in the bead engulfment assay. To be sure, that beads outside the cells were not counted, stack images were checked and snaps with least number of beads (3-5) lying outside the cells were chosen for further analysis so as to get cleaner, noise free and unbiased counting of beads engulfed by the cells. For each biological replicate, multiple cells (∼ 10 – 15) were analyzed using the aforementioned analysis per coverslip, for a total of 6 coverslips per biological replicate. Based on this analysis, the number of beads engulfed per cell for each biological replicate was averaged across all the cells counted for that biological replicate, and graphically plotted as a single averaged value per biological replicate.

### Gel based ABPP assays

Microglial cells were harvested by gently scraping in chilled PBS, lysed using probe sonicator (1 sec on, 3 secs off, 60% amplitude, 30 seconds) on ice, the membrane lysates were prepared as per protocols reported by us previously^*10, 22, 27*^, and the protein concentration in these membrane lysates was measured using the Pierce BCA kit as per manufacturer’s instructions. The membrane lysates were diluted to a final concentration of 1 mg/mL for all the gel-based ABPP assays. For competitive ABPP, these membrane lysates were first incubated with 10 μM JJH350 or vehicle (DMSO) at 37 °C for 45 mins with constant shaking in a final volume of 100 μL. Thereafter, samples were incubated with fluorophosphonate-rhodamine (FP-Rh) probe (2 μM) at 37 °C for 45 mins with constant shaking recently reported by us^*10, 28, 32*^. Finally, the reactions were quenched by adding 40 μL of 4X-SDS-PAGE loading dye and boiling the samples for 10 mins at 95 °C. For Cu-based alkyne azide click (CuAAC) chemistry based JJH350 labelling reactions, 100 μL membrane lysates (at 1 mg/mL) were treated with 10 μM JJH350 or vehicle (DMSO) at 37 °C for 45 mins with constant shaking Thereafter, these treated membrane lysates were treated with click mixture for 45 mins at 37 °C with constant shaking. The click mixture (11 μL per reaction of 100 μL lysate) consisted of 6 μL of 1.7 mM TBTA in 4:1 DMSO: *tert* butanol, 2 μL of 50 mM copper sulfate, 2 μL of 50 mM TCEP and 1 μL of rhodamine-azide (10 mM in DMSO) as per standard protocols reported earlier^*31, 33*^. The click chemistry reactions were quenched by adding 45 μL of 4X SDS-PAGE loading dye. All the gel-based ABPP samples were resolved on a 10% SDS-PAGE gel and imaged for in-gel fluorescence using the Syngene G-Box Chemi-XRQ gel documentation system as per standard protocols reported by us^*28, 32*^.

### Statistical analysis

All data in this paper was plotted and statistically analyzed using the Prism 9 (version 9.1.0) software for macOS (GraphPad). All bar data in this paper is represented as the mean ± standard deviation (SD) from at least three independent biological replicates per experimental group. The Student’s two-tailed unpaired parametric *t*-test was used to determine the statistical significance between different experimental groups, and a *p*-value of < 0.05 was considered statistically significant in this study.

## RESULTS

### ABHD12-null mice show heightened phagocytosis in the cerebellum

The loss of ABHD12 activity in mice results in an accumulation of brain lyso-PSs, particularly VLC lyso-PSs^*15, 24*^, that over time, causes microglial activation, especially in the arbor vitae region closer to the deep nuclei in the cerebellum, given the localization of ABHD12 and the enrichment of the upstream PS lipase (major lyso-PS biosynthetic enzyme) ABHD16A ^*27*^ to this anatomical region, and therefore, results in neuroinflammation^*24*^. Following up on these seminal findings, we have recently shown from studies in the mammalian (mouse) brain, that during inflammation caused by oxidative stress, ABHD12 also controls the levels of the potent pro-apoptotic oxidized PSs, and its deletion results in increased brain oxidized PS levels (and likely manifests into neuroinflammation)^*10*^. Given these recently published studies and the strong association of lyso-PS/oxidized PS and ABHD12 with neuroinflammation (particularly in the cerebellum), we wanted to explore the effects of deletion of ABHD12 on microglial activation during an inflammatory stimulus. In our current studies, this inflammatory stimulus was provided by systemically injecting mice with lipopolysaccharide (LPS) (10 mg/kg body weight, intraperitoneally, 4 hours), following which, the brains of these treated mice were isolated, processed for immunohistochemical analysis and the extent of microglial activation was assessed by counting the number of ionized calcium binding adaptor molecule (Iba-1) stained cells having their soma size (area) > 200 μm^2^ per unit area in arbor vitae region of the cerebellum, using protocols recently reported by us^*15, 27*^. From these studies, we found that, while the inflammatory stimulus (LPS injection) markedly increased the number of activated microglia in the cerebellum (∼ 10-fold) relative to the vehicle (1x-phosphate buffered saline, PBS) control, we did not notice any significant differences in total numbers between the two different genotypes i.e., the wild type (+/+) and ABHD12-null (–/–) mice groups, for either the vehicle or LPS treatments (**Figure 2A**).

**Figure 2.**
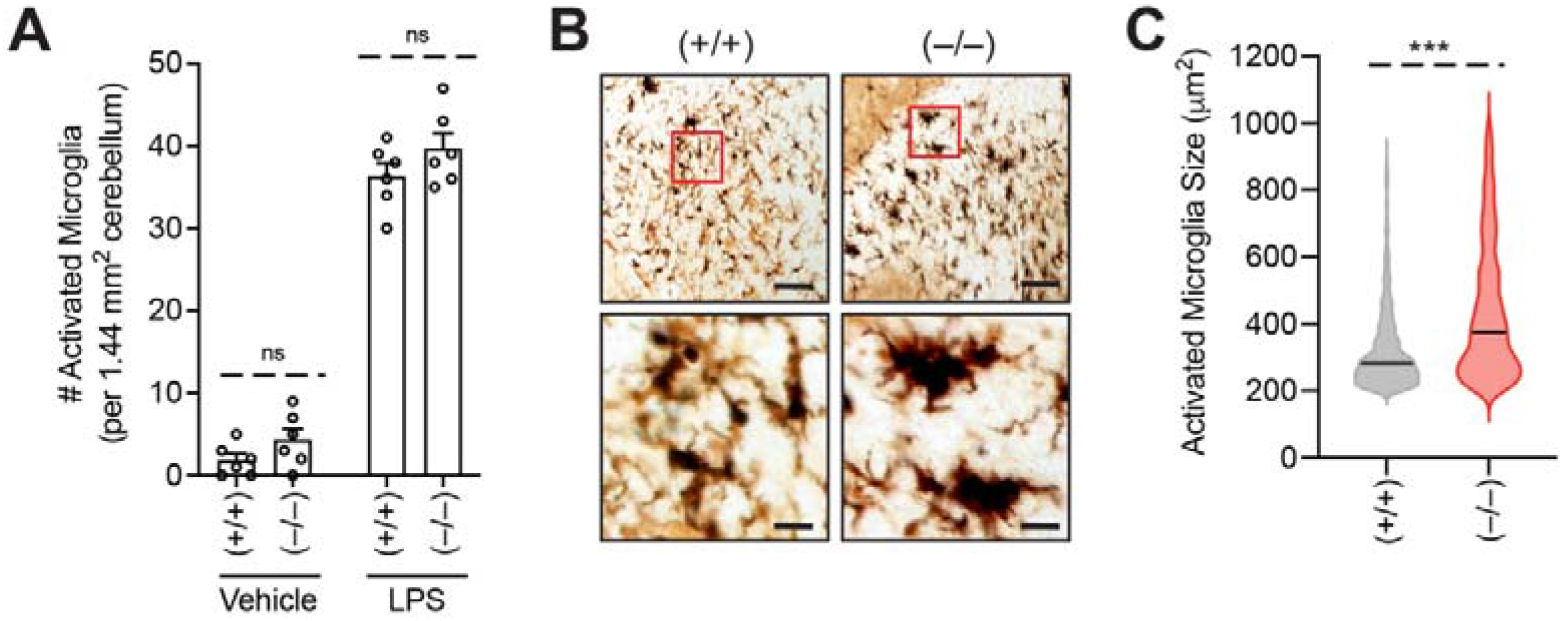
ABHD12-null activated microglia in the arbor vitae region of the cerebellum have increased size. (**A**) Quantification of activated microglia (enlarged cells, > 200 μm^2^ area) in the arbor vitae region of the cerebellum (per 1.44 mm^2^ area) of wild type (+/+) and ABHD12 knockout (–/–) mice following intraperitoneal injection of vehicle (1x PBS) or LPS (10 mg/kg body weight) for 4 hours. Data represents mean ± standard deviation from six independent experiments (biological replicates) per experimental group. (**B**) Representative Iba-1 immunostaining image of the activated microglia in the arbor vitae region of the cerebellum of wild type (+/+) and ABHD12 knockout (–/–) mice following intraperitoneal injection of LPS (10 mg/kg body weight, 4 hours), showing amoeboid morphology for activated microglia from the ABHD12 knockout mice. This experiment was done 6 times with reproducible results each time. Top images, scale bar = 250 μm; bottom zoomed images, scale bar = 10 μm. (**C**) Size of activated microglia in the arbor vitae region of the cerebellum of wild type (+/+) and ABHD12 knockout (–/–) mice following intraperitoneal injection of LPS (10 mg/kg body weight, 4 hours), showing increased average size of activated microglia from the ABHD12 knockout mice. Violin plot data represents ensemble of sizes of 1085 individual activated microglia from the cerebellum from six independent experiments (biological replicates) per experimental group, and the black line in the plot, represents average size of activated microglia for that experimental group. ***p < 0.001 versus (+/+) group by Student’s two-tailed unpaired parametric *t*-test; ns = not statistically significant.

Interestingly, from this immunohistochemical experiment, we visually noticed that upon the inflammatory stimuli, activated microglia from both genotypes had different morphologies (**Figure 2B**). Upon closer inspection, we found that following the inflammatory stimulus, the activated microglia from wild type (+/+) mice had significantly more ramifications, while those from ABHD12-null (–/–) mice adopted amoeboid shapes (**Figure 2B**), consistent with a heightened microglial activation state for the latter^*34-36*^. To ensure that there were no observational biases in our visual inspections, we performed these experiments blinded, and computed the average size (area in μm^2^) of the activated microglia in the arbor vitae region close to the deep cerebellar nuclei in the brains of mice from both genotypes upon the inflammatory stimuli using the automated image analyzing tool, ImageJ^*29, 37*^. Here, we found that consistent with our visual observation of the amoeboid morphology, from six independent experiments (biological replicates), the average size of activated microglia was significantly larger in ABHD12 knockout mice (**Figure S1**). Next, we pooled all the data for size of the activated microglia for both genotypes from these six biological replicates (N = 1085), and found that the activated microglia from ABHD12-null (–/–) mice had significantly greater average size (∼ 434 μm^2^) compared to those from wild type (+/+) mice (∼ 325 μm^2^) (**Figure 2C**).

Heightened activation and an amoeboid morphology upon an inflammatory stimulus are both directly associated with increased phagocytosis activity by the microglia^*34-36*^. Previously, we have also shown that during the same inflammatory stimulus, there is an increased production of pro-inflammatory cytokines (TNF-α and IL-6) in the brains of ABHD12 knockout mice relative to wild type controls, consistent with an increased activation status of the microglia of ABHD12-null mice^*10*^. Given this, we next wanted to investigate, if this was indeed the case, and if this was consistent with our immunohistological observations of activated microglia in the arbor vitae region of the cerebellum of ABHD12-null (–/–) mice. Towards this, using established immunohistochemical analysis^*15, 27*^, we checked the activated microglia, for the presence of the lysosome-associated membrane protein 1 (LAMP1), as this protein is present on late-stage phagosomes^*38, 39*^, and serves as a good indicator phagocytotic activity by macrophages and/or microglia. From this experiment, which were performed blinded, we found that upon an inflammatory stimulus, activated cerebellar microglia from ABHD12-null (–/–) mice had significantly more LAMP1 levels (∼ 2-fold) relative to those from wild type (+/+) mice (**Figure 3**), consistent with an increased phagocytosis activity in the former, along with our previous finding^*10*^, of the increased pro-inflammatory cytokine production in the brains of ABHD12 knockout mice upon the same inflammatory stimulus.

**Figure 3.**
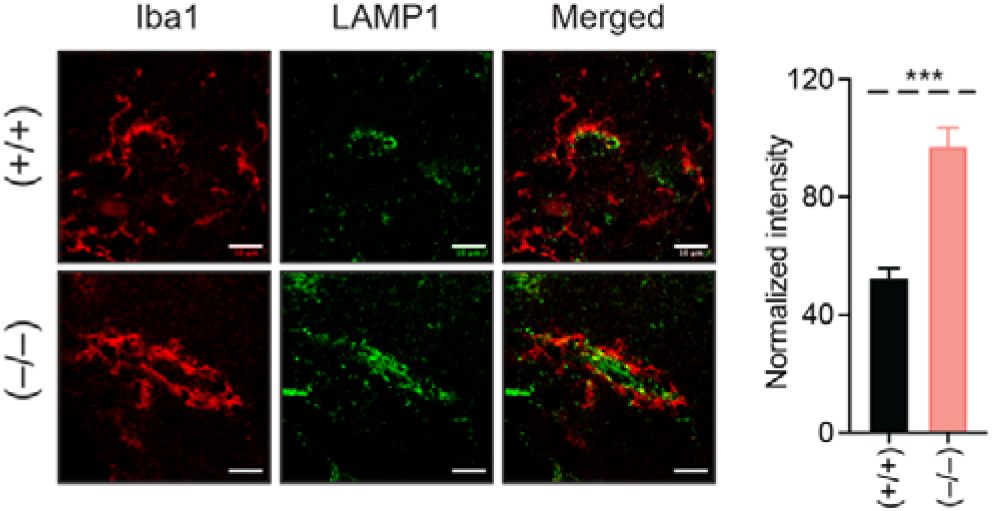
ABHD12-null activated microglia in the cerebellum have increased LAMP1 expression. Representative immunofluorescence images and quantification of LAMP1 levels on activated microglia in the cerebellum of wild type (+/+) and ABHD12 knockout (–/–) mice following intraperitoneal injection of LPS (10 mg/kg body weight, 4 hours), showing a ∼ 2-fold increase in the levels of LAMP1 on activated microglia from ABHD12 knockout mice. Data (bar plots) is represented as mean ± standard deviation from six independent experiments (biological replicates) per experimental group. Scale bar in the immunofluorescence images is 5 μm. ***p < 0.001 versus (+/+) group by Student’s two-tailed unpaired parametric *t*-test.

### Brain ABHD12 levels increase upon inflammatory stimuli

Previous studies in primary macrophages have shown that upon an inflammatory stimulus (LPS treatment), the levels and in turn the lipase activity of ABHD12 significantly decreases (∼ 2-fold) with a concomitant increase in the concentrations of the lipid metabolites that it regulates i.e. the cellular oxidized PS lipids and secreted lyso-PS lipids^*10, 23, 27*^. In addition to this, we have previously shown in a LPS-based neuroinflammation paradigm, that the concentration of lyso-PSs in the brains of mice remain unchanged, while those of oxidized PS significantly increase, and that ABHD12 regulates lyso-PS levels during basal conditions (vehicle treatments), and oxidized PS levels during oxidative stress (LPS treatment)^*10*^. Given this precedence, we wanted to know if the levels of ABHD12 in the wild type mammalian (mouse) brain indeed change during an inflammatory stimulus (LPS injection, 10 mg/kg body weight, intraperitoneally, 4 hours). We decided to investigate this initially using immunohistochemical analysis of microglial cells in the arbor vitae region closer to the deep cerebellar nuclei, as previous studies have shown that primary microglia have high protein levels and hence increased activity of ABHD12^*40*^. Quite surprisingly, and opposite to the findings from primary macrophages^*23, 27*^, we found that upon an inflammatory stimulus, the concentration of ABHD12 on activated microglia of the cerebellum actually significantly increased (∼ 2-fold) (**Figure 4A**). To further validate this finding, we decided to perform an immunoblotting (western blot) experiment on membrane lysates of the cerebellum, following an inflammatory stimulus. Here, we found consistent with the immunohistochemical analysis (**Figure 4A**), that relative to the vehicle (PBS) control, the levels of ABHD12 in the cerebellum were indeed significantly elevated (∼ 2-fold) upon an inflammatory stimulus (LPS-injection, 10 mg/kg body weight, intraperitoneally, 4 hours) (**Figure 4B**). In this immunoblotting experiment, the cytoskeleton protein β-actin was used as an equal loading control (**Figure 4B**). In primary macrophages, it has also been shown that the protein levels and therefore the lipase activity of the major lyso-PS biosynthetic enzyme (PS lipase) ABHD16A significantly increases upon an inflammatory stimulus^*23, 27*^, and hence we decided to check if the levels of this PS-lipase were altered in our immunoblotting experiment on membrane lysates of the cerebellum. Interestingly, we did not find any changes in levels of the PS lipase ABHD16A in the cerebellum upon an inflammatory stimulus (**Figure 4B**), consistent with our previous findings, that show that the lyso-PS levels remain unchanged in a LPS-based neuroinflammation paradigm^*10*^.

**Figure 4.**
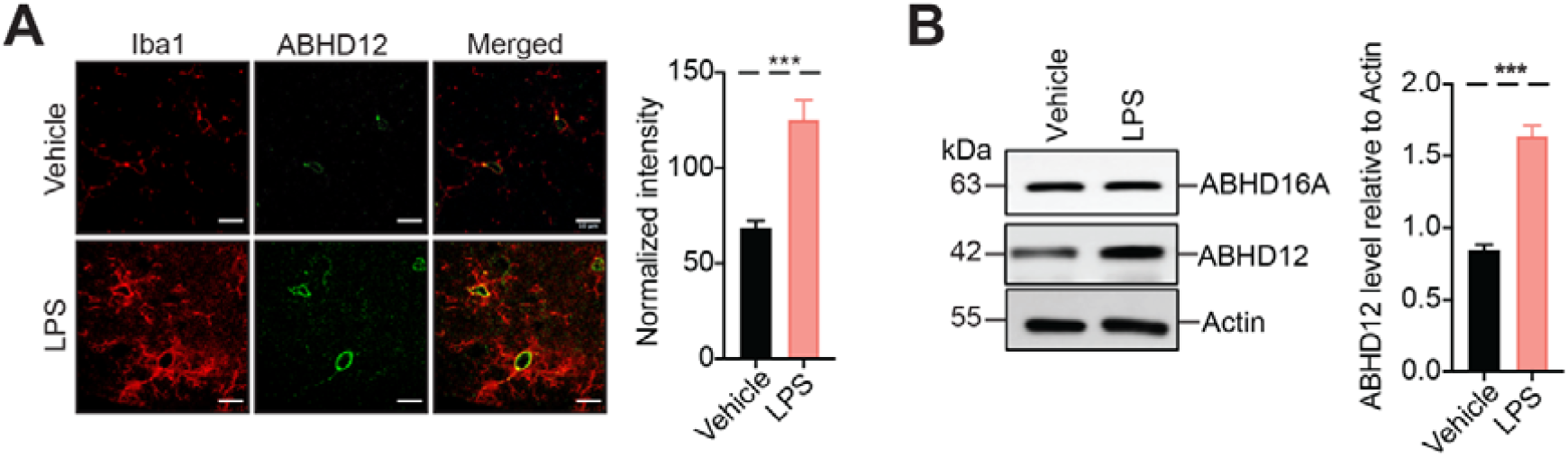
Brain ABHD12 levels increase upon an inflammatory stimulus. (**A**) Representative immunofluorescence images and quantification of ABHD12 levels on cerebellar microglia from wild type (+/+) mice, following intraperitoneal injection of vehicle (1x PBS) or LPS (10 mg/kg body weight, 4 hours), showing a ∼ 2-fold increase in ABHD12 levels following LPS treatment. Data (bar plots) is represented as mean ± standard deviation from six independent experiments (biological replicates) per experimental group. Scale bar in the immunofluorescence images is 10 μm. (**B**) Representative blot and quantification from Western blot analysis of membrane lysates of the cerebellum from wild type (+/+) mice, following intraperitoneal injection of vehicle (1x PBS) or LPS (10 mg/kg body weight, 4 hours), showing a ∼ 2-fold increase in ABHD12 levels following LPS treatment. The levels of the PS lipase ABHD16A in the cerebellum did not change following LPS treatment. Actin was used as a loading control for this experiment. This experiment was done three times (3 independent biological replicates) with reproducible results each time. For both (**A**) and (**B**), ***p < 0.001 versus (+/+) group by Student’s two-tailed unpaired parametric *t*-test.

### Disruption of ABHD12 increases phagocytosis by microglial cells

That upon an inflammatory stimulus, the microglia in ABHD12-null mice have heightened activation status, an amoeboid morphology, and higher levels of the phagocytic marker LAMP1 strongly points to a role of ABHD12 in modulating phagocytosis (**Figure 2, 3**). Additionally, the increase in cerebellar ABHD12 protein levels upon an inflammatory stimulus, further suggests that this lipase controls levels of the pro-apoptotic oxidized PS in the mammalian brain during oxidative stress, and possibly tempers heightened phagocytosis during such physiological conditions (**Figure 4**). Given these findings, we next wanted to assess, whether the pharmacological inhibition and/or genetic deletion of ABHD12 in microglial cells had any effect on the rate of phagocytosis or phagocytic activity in these innate immune cells. For these experiments, we first chose the well-studied BV-2 and N9 immortalized microglial cell lines, and first confirmed that these microglial cells indeed express ABHD12 by western blotting (**Figure S2**). Next, we also confirmed by established competitive gel-based activity based protein profiling (ABPP) assays^*10, 41, 42*^, that the *N*-hydroxyhydantoin-carbamate JJH350 (10 μM, 6 hours), a recently reported ABHD12 selective covalent inhibitor^*31*^, indeed reacted with and covalently inhibited ABHD12 in an irreversible manner in both the BV-2 and N9 microglial cells, and was therefore used for the pharmacological inhibition of ABHD12 in all our subsequent assays in these cell lines (**Figure S3**). Next, to determine the effect of ABHD12 inhibition on phagocytosis, we treated the BV-2 or N9 cells with vehicle (DMSO) or JJH350 (10 μM, 6 hours), following which, they were incubated with fluorescent latex beads (0.5 μm, cell to bead ratio = 1:200 for 30 mins), and the number of beads that were engulfed by the microglial cells was visualized by fluorescence microscopy using established protocols^*27, 28*^. It is known that these fluorescent latex beads act as antigens, and are known to trigger oxidative stress response due to activation of inflammatory and/or phagocytic pathways in phagocytes^*43, 44*^. From these bead-engulfment (phagocytosis) assays, we found that majority of the microglial cells (∼ 90% of the total cells) engulfed the fluorescent latex beads, and for either treatment (vehicle or JJH350), the percentage of total cells engulfing the fluorescent latex beads for both the BV-2 and N9 microglia was almost identical (**Figure 5A**). On closer visual inspection of the microscopy images, we found that the JJH350 treated microglia engulfed markedly more fluorescent beads than the vehicle treated microglia for both the BV-2 and N9 microglial cells (**Figure 5B**). Next, upon counting the number of fluorescent latex beads engulfed per phagocytosing microglial cell, we found that the JJH350 treated microglia engulfed ∼ 3-times more fluorescent latex beads than the corresponding vehicle treated microglia (average fluorescent latex beads engulfed per phagocytosing cell for BV-2 microglia: vehicle = 12, JJH350 = 33; N9 microglia: vehicle = 11, JJH350 = 30) (**Figure 5C**).

**Figure 5.**
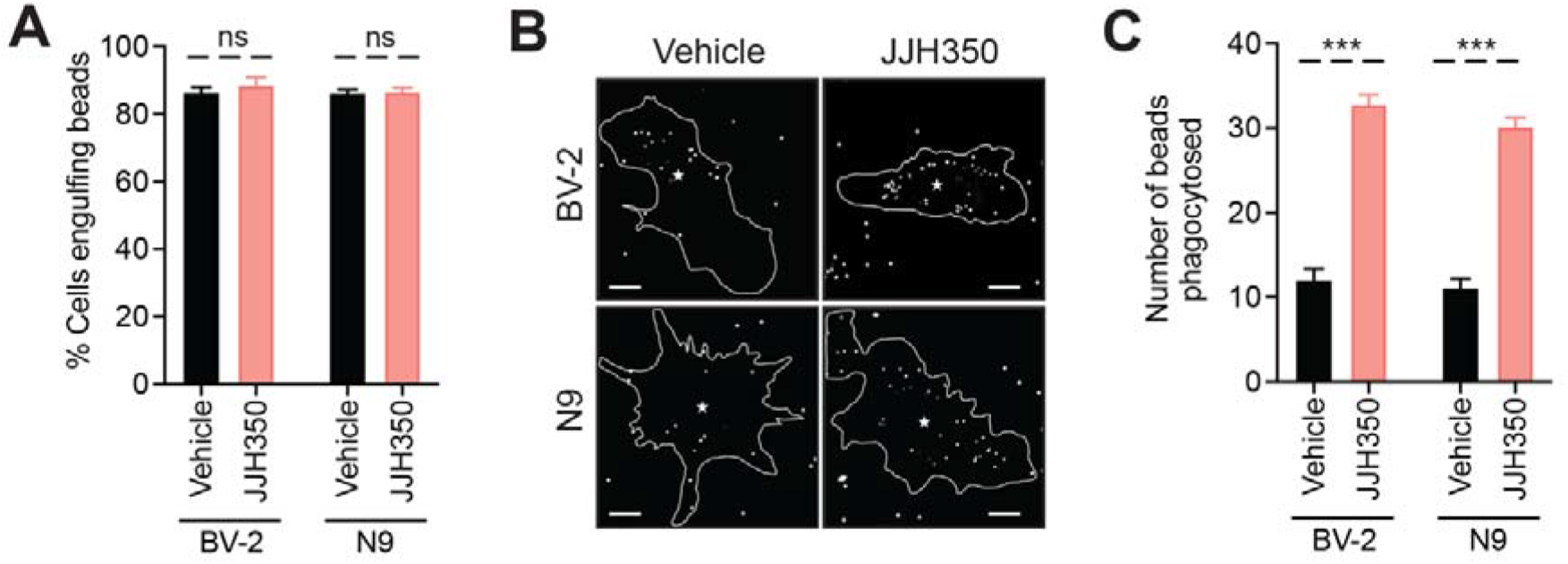
Pharmacological inhibition of ABHD12 results in increased phagocytosis by microglial cells in a fluorescent bead uptake assay. (**A**) Quantification of the total percentage of microglial cells (BV-2 and N9) engulfing fluorescent beads following vehicle (DMSO) or JJH350 (10 μM, 6 hours) treatment, showing no significant differences between the two experimental groups. Data (bar plots) is represented as mean ± standard deviation from six independent experiments (biological replicates) per experimental group. (**B**) Representative microscopy images of microglial cells (BV-2 and N9) engulfing fluorescent beads following vehicle (DMSO) or JJH350 (10 μM, 6 hours) treatment, showing increased phagocytosis of fluorescent beads in the JJH350-treated microglial cells. Scale bar in the microscopy images is 10 μm and (*) represents centroid of the nucleus. (**C**) The number of fluorescent beads phagocytosed by microglial cells (BV-2 and N9) following vehicle (DMSO) or JJH350 (10 μM, 6 hours) treatment, showing a significant increase of number of phagocytosed fluorescent beads by the JJH350-treated microglial cells (BV-2 and N9). Data (bar plots) is represented as mean ± standard deviation from six independent experiments (biological replicates) per experimental group. ***p < 0.001 versus vehicle group by Student’s two-tailed unpaired parametric *t*-test; ns = not statistically significant.

To complement these pharmacological studies, using established protocols^*40*^, we harvested primary microglial cells from wild type (+/+) and ABHD12 knockout (–/–) mice, and performed the same fluorescent latex bead engulfment assays to determine, whether the genetic deletion of ABHD12 also has any effect on the phagocytic activity of primary microglial cells. In these cellular phagocytosis assays, we found that the majority of the primary microglial cells (∼ 90% of the total cells) engulfed the fluorescent latex beads, and in terms of the percentage of total primary microglial cells engulfing these fluorescent beads, we found that for both the genotypes these numbers were almost identical (**Figure 6A**). Upon closer visual inspection of the microscopy images from these cellular phagocytosis assays, we found that the ABHD12-null (–/–) primary microglial cells engulfed significantly more fluorescent beads compared to wild type (+/+) primary microglial cells (**Figure 6B**). On quantifying the number of fluorescent latex beads engulfed per phagocytosing primary microglial cell, we found that the ABHD12-null (–/–) primary microglial cells engulfed ∼ 2-times more fluorescent beads than the corresponding wild type (+/+) primary microglia (average fluorescent latex beads engulfed per phagocytosing cell by primary microglia: wild type (+/+) = 65, and ABHD12-null (–/–) = 150) (**Figure 6C**). Interestingly, we also performed these cellular phagocytosis assays in primary thioglycollate-elicited peritoneal macrophages derived from wild type (+/+) or ABHD12-null (–/–) mice^*10, 27*^, and found results similar to those observed in the microglial cells, that the loss of ABHD12 activity in macrophages resulted in significantly more (∼ 3-times) phagocytosis of fluorescent latex beads (**Figure S4**).

**Figure 6.**
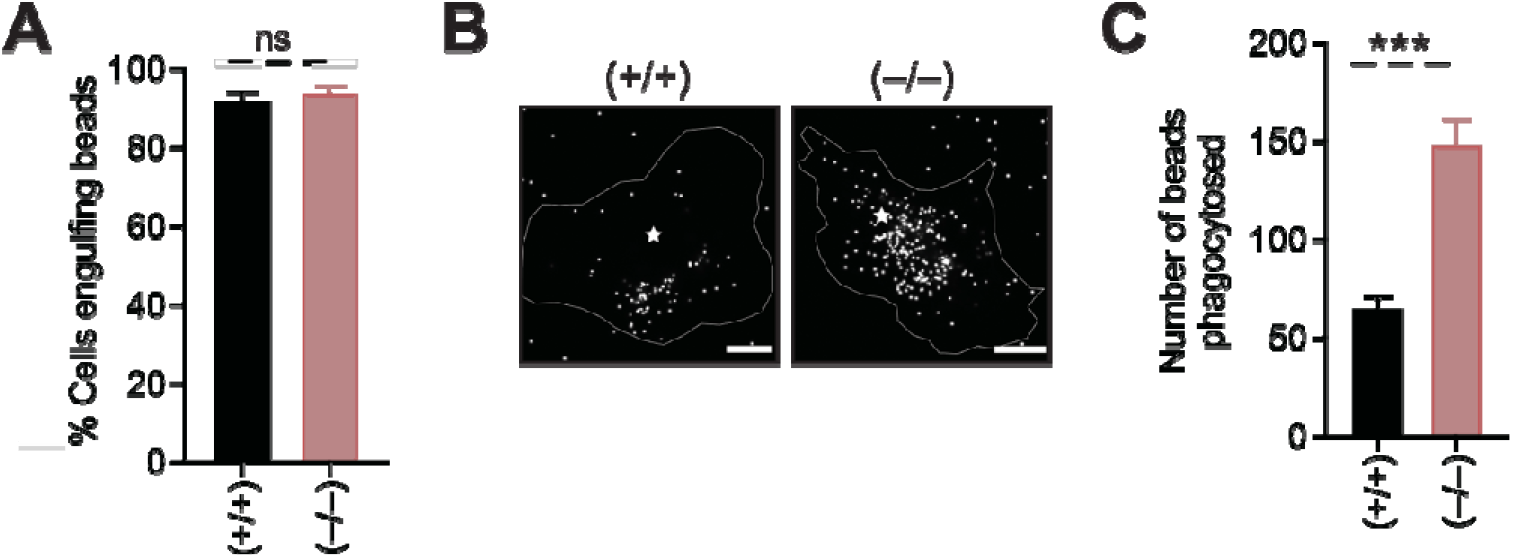
Deletion of ABHD12 in primary microglial cells results in increased phagocytosis in a fluorescent bead uptake assay. (**A**) Quantification of the total percentage of primary microglial cells obtained from wild type (+/+) and ABHD12 knockout (–/–) mice engulfing fluorescent beads, showing no significant differences between the two genotypes. Data (bar plots) is represented as mean ± standard deviation from six independent experiments (biological replicates) per genotype. (**B**) Representative microscopy images of primary microglial cells obtained from wild type (+/+) and ABHD12 knockout (–/–) mice, engulfing fluorescent latex beads, showing increased phagocytosis activity towards engulfing the fluorescent latex beads in the ABHD12-null primary microglial cells. Scale bar in the microscopy images is 10 μm and (*) represents centroid of the nucleus. (**C**) The number of fluorescent beads phagocytosed by primary microglial cells obtained from wild type (+/+) and ABHD12 knockout (–/–) mice, showing a significant increase of number of phagocytosed fluorescent latex beads by ABHD12-null primary microglial cells. Data (bar plots) is represented as mean ± standard deviation from six independent experiments (biological replicates) per genotype. ***p < 0.001 versus vehicle group by Student’s two-tailed unpaired parametric *t*-test; ns = not statistically significant.

## DISCUSSION

Little over a decade ago, deleterious null mutations in the gene that encodes the integral membrane-associated lipase ABHD12 was identified as the cause for the early onset human neurological disorder PHARC, which is clinically represented in human subjects as polymodal sensory and motor defects caused by peripheral neuropathy, coupled with an early onset of cataract (and/or blindness), and hearing loss. Since this initial discovery, several studies have been published, that describe the biochemical activities of ABHD12 and the lipid signaling pathways that this disease-associated lipase regulates in mammals, including humans. In a seminal study by Cravatt and co-workers, they generated the ABHD12 knockout mouse, and showed this rodent model serves as an excellent pre-clinical surrogate to study various aspects of the pathophysiology of PHARC. In the same study, it was chronologically shown that the loss of the ABHD12 lipase activity first results in the accumulation of the bioactive lyso-PSs in the brain (**Figure 1**), that causes deregulated (lyso-PS-mediated) signaling over time, particularly in the cerebellum (given the localization of ABHD12 and the upstream lyso-PS biosynthetic enzyme ABHD16A in the cerebellum) of ABHD12 knockout mice. This aberrant lyso-PS signaling is then succeeded by microglial activation in the arbor vitae region of the cerebellum and thereby (neuro)inflammation, which likely manifests into behavioral (motor and auditory) defects in the ABHD12 knockout mice, mirroring those seen in human PHARC subjects (**Figure 1**). Following up on these findings, we have recently shown that, by its lipase activity, ABHD12 also controls the physiological concentrations of the potent pro-apoptotic oxidized PS lipids (**Figure 1**), and significantly tempers inflammatory responses under conditions of oxidative stress, particularly in the mammalian brain and macrophages. Interestingly, both the lyso-PS and oxidized PS lipids have had a strong association with the activation of macrophage cells, and phagocytosis, and yet despite its biomedical importance, the effect that the loss of ABHD12 activity has on phagocytosis and the correlation of phagocytosis to the pathological phenotypes associated with PHARC, has not been thoroughly investigated or reported to the best of our knowledge.

Given the role of ABHD12 in modulating levels of the immunomodulatory lyso-PSs and pro-apoptotic oxidized PSs in macrophages, we decided to investigate whether the loss of its lipase activity has any effect on phagocytosis. Towards this, we first challenged wild type and ABHD12 knockout mice with an inflammatory stimulus (systemic LPS injection), and checked the extent of neuroinflammation in the arbor vitae region of the cerebellum, by counting the number of activated microglia using established immunohistochemical analysis. We found in this anatomical region of the cerebellum, that while the total number of activated microglia between the two genotypes did not change, those from ABHD12 knockout mice adopted an amoeboid morphology, and had a greater soma size (**Figure 2**), with a concomitant increase in the phagocytic marker protein LAMP1 (**Figure 3**), all together consistent with a heightened phagocytosis activity in ABHD12-null cerebellar microglia. We have previously shown that the pro-apoptotic oxidized PS lipids accumulate in the brains of ABHD12 knockout mice during conditions of oxidative stress (e.g. during systemic LPS injection). Interestingly, and contrary to findings in macrophages, here we show that during such oxidative stress conditions, in the wild type mice brains, the levels of ABHD12 increase nearly two-fold (**Figure 4**). We think this is an innate anticipatory mechanism in the mammalian brain, where by increasing the levels of ABHD12 in response to an inflammatory stimulus, excess oxidized PSs produced during oxidative stress in the brain (particularly in the cerebellum) are efficiently metabolized by the oxidized PS lipase activity of ABHD12, and therefore the apoptosis and/or phagocytosis due to the accumulation of the oxidized PS lipids in neurons or other brain resident glial cells is prevented. Next, we wanted to directly assess in cultured microglial cells, if the loss of ABHD12 activity indeed resulted in increased phagocytosis, and decided to address this directly using an established fluorescent latex bead-uptake assay using complementary pharmacological and genetic approaches. We found from this assay in two immortalized microglial cell lines (BV-2 and N9), that the pharmacological inhibition of ABHD12 activity resulted in significantly more phagocytosis activity, and hence an increased uptake of fluorescent latex beads in both cell lines (**Figure 5**). We also performed the same bead-uptake assay in primary microglia cells that obtained from wild type (+/+) or ABHD12-null (–/–) mice, and found that the deletion of ABHD12 and loss of its lipase activity in primary microglial cells results in significantly more uptake of fluorescent latex beads via phagocytosis (**Figure 6**). Interestingly, we found that the loss of the ABHD12 activity also resulted in similar increased phagocytosis activity in primary peritoneal macrophages (**Figure S4**). Taken together, all our data, provide compelling evidence both *in vivo* and by *in vitro* biochemical experiments, that the loss of ABHD12 activity indeed causes heightened phagocytosis activity in microglial (and macrophage) cells.

We think that the association of loss of ABHD12 activity with increased phagocytosis activity has tremendous implications in the context of the pathophysiology of PHARC, and propose a model based on the findings in this paper, for explaining this (**Figure 7**). We have recently shown why the cerebellum is the most atrophic region of the brain in human PHARC subjects given the distinct localization of ABHD12 and ABHD16A to the Purkinje and Granular neurons respectively^*27*^. During normal conditions, the microglia are constantly palpating various neurons present in the cerebellum, and phagocytosis by these microglia is at a minimal, given that most cerebellar neurons are healthy, and signaling events are fairly homeostatic and well-regulated at this point. In human PHARC subjects, we speculate that the loss of ABHD12 activity results in the buildup lyso-PS (even during basal conditions because of the sustained PS lipase activity of ABHD16A in the cerebellum)^*23, 24, 27*^ and oxidized PS (during some inflammatory stimuli, oxidative stress)^*10*^ in the cerebellum, presumably from Purkinje neurons and cerebellar microglial cells that highly express ABHD12 in this brain region. Together, the increased concentrations of the immunomodulatory lyso-PS and pro-apoptotic oxidized PS lipids because of loss of ABHD12 activity, further activate cerebellar microglia, and initiate a heightened phagocytosis response, that we posit, leads to the continued neuronal damage in the cerebellum. This damage to cerebellar (Purkinje) neurons eventually manifests into the sensory and motor defects observed in human PHARC subjects. We also speculate that the heightened phagocytosis activity by microglial cells due to the loss of ABHD12, over time, also causes the cerebellar atrophy observed in human PHARC subjects.

**Figure 7.**
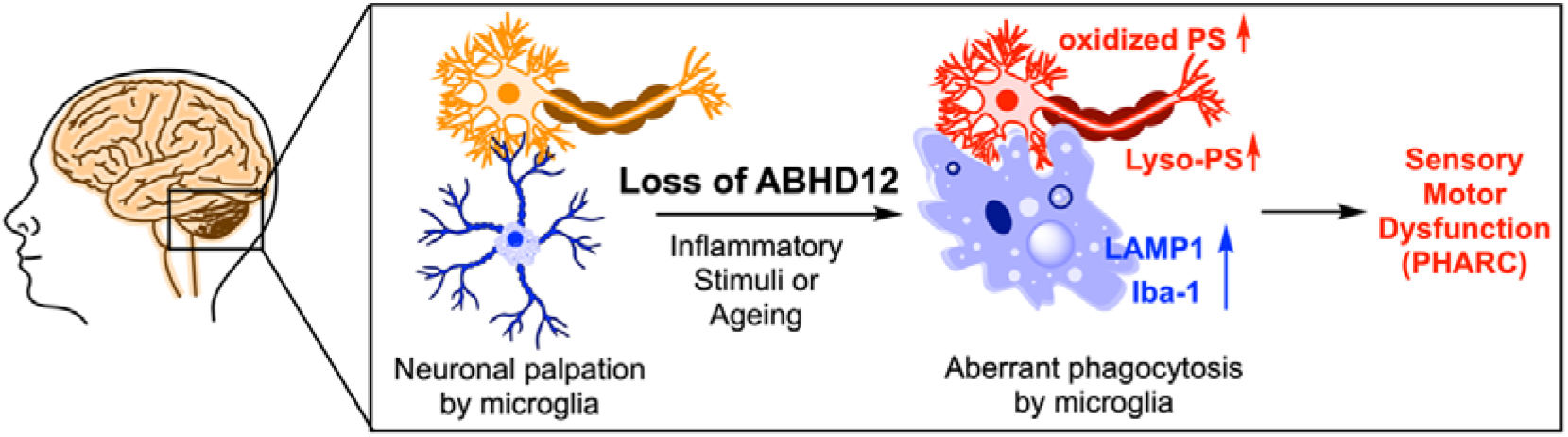
A model summarizing the increased phagocytosis observed in the cerebellar microglial cells in ABHD12 knockout mice.

Moving ahead, it will be interesting to mechanistically understand and delineate how the flux and interplay between lyso-PSs and oxidized PSs intricately regulated by ABHD12 in the brain, particularly in the cerebellar microglial cells and neurons, influence the various phagocytosis pathways that culminate into neuroinflammation, and lead to the behavioral-defects associated with PHARC. Towards understanding this, it will be very useful to generate much needed brain cell-specific ABHD12 knockout mice models (e.g. deletion of ABHD12 specifically in neurons or microglia) to map this interplay in the central nervous system, and dissect out contribution of different cells in the brain in causing the pathology observed in PHARC. Currently, very little (if at all anything) is known of the putative receptors for both lyso-PSs and oxidized PSs in microglial cells and the mammalian brain in general, and therefore identifying and annotating such as-of-yet unknown receptors, will add tremendous value to our understanding of phagocytosis pathways in the brain associated with both these signaling lipids. We have recently shown that VLC lyso-PSs, that accumulate most in the brains of ABHD12 knockout mice, potently activate macrophages via a TLR2-dependent pathway^*15*^, and it would be interesting to mechanistically investigate the VLC lyso-PS – TLR2 axis in the context of phagocytosis and neuroinflammation observed in PHARC. Finally, on a same note, from a translational perspective, it will also be interesting to assess if the microglial activation associated with ABHD12 deletion in the murine model of PHARC, can be rectified by treatment with pharmacological agents such as the neuroprotective antibiotic minocycline, and if so, this can be translated into a therapy for treating the neuroinflammation observed in human PHARC subjects.

## Supporting information

Suppplementary Figure 1 - 4

## ASSOCIATED CONTENT

### SUPPORTING INFORMATION

The Supporting Information is available free of charge at xxxx. Figures S1 – S4 (PDF).

### ACCESSION CODES

The Uniprot IDs for mouse and human ABHD12 are Q99LR1 and Q8N2K0 respectively.

### AUTHOR CONTRIBUTIONS

S.S. performed all the experiments. S.S. and S.S.K. conceived the project, analyzed all the data and wrote the manuscript. S.S.K. acquired funding and supervised this study.

### FUNDING

This work was supported by funding from a DBT/Wellcome Trust India Alliance Fellowship (grant number: IA/I/15/2/502058) and Science and Engineering Research Board (SERB) (grant number: CRG/2020/000023) to S.S.K. Part of this work was carried at the National Facility for Gene Function in Health and Disease (NFGFHD) at IISER Pune, which is supported by a grant from the Department of Biotechnology, Govt. of India (BT/INF/22/SP17358/2016). S.S. is supported by a graduate student fellowship from IISER Pune.

### NOTES

The authors declare no competing financial interests.

## ACKNOWLEDGEMENTS

The IISER Pune – Leica Microscopy facility is thanked for providing access to the microscopes needed for the imaging studies described in this paper.

## TABLE OF CONTENT GRAPHICS

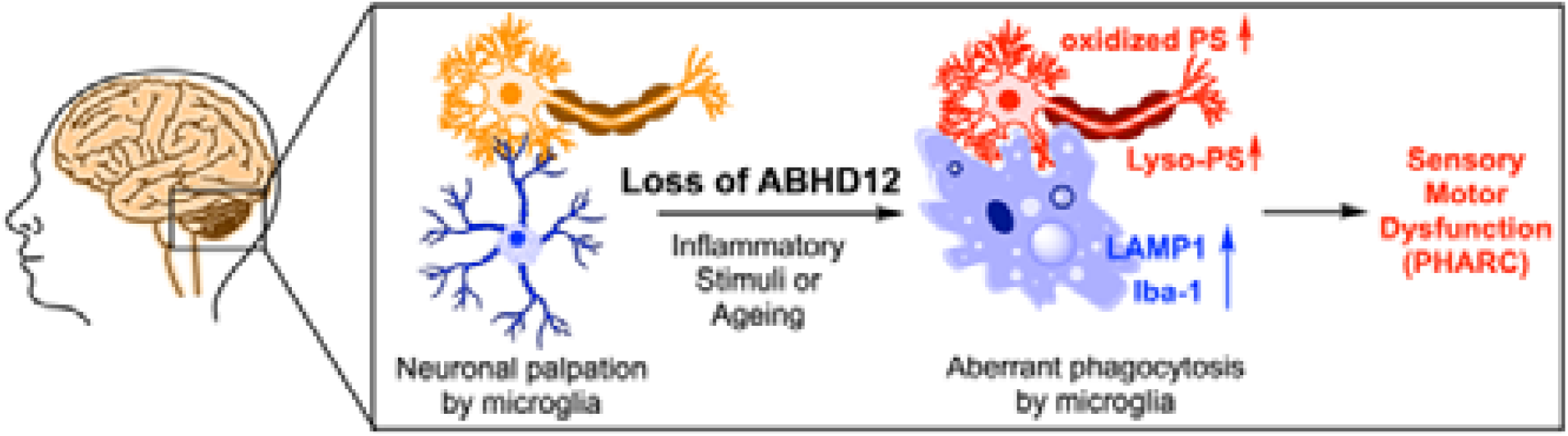

