## Supplementary material for "The loss of enzymatic activity of the PHARC associated lipase ABHD12 results in increased phagocytosis that causes neuroinflammation": Suppplementary Figure 1 - 4

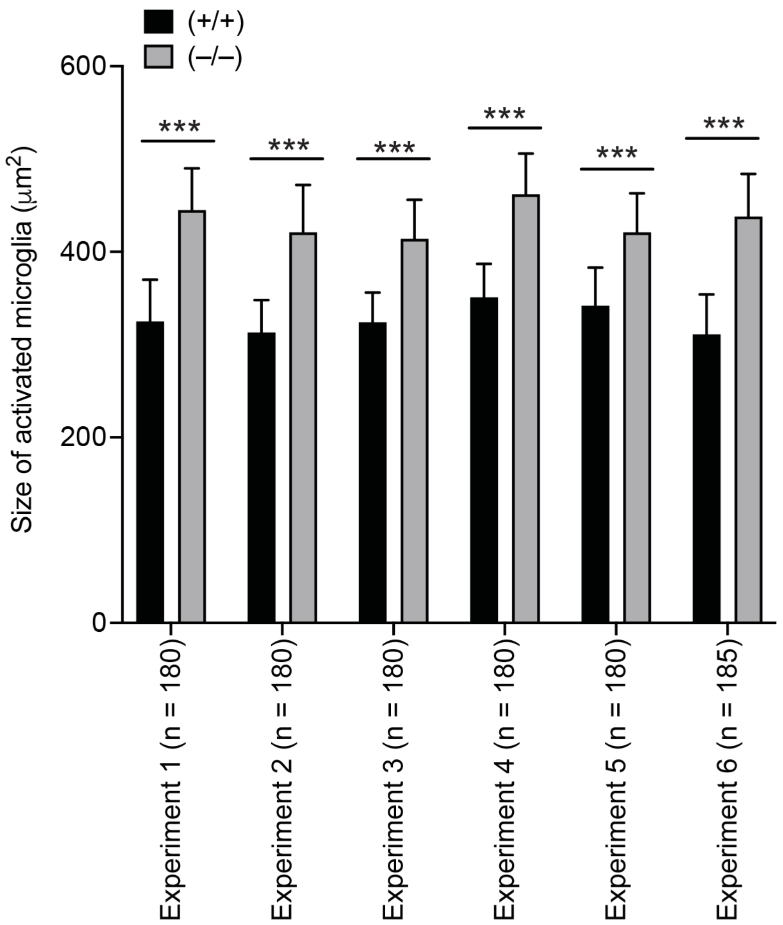


**Figure S1**. **Size of activated cerebellar microglial cells per biological replicate.** The size of activated microglia in the arbor vitae region of the cerebellum of wild type (+/+) and ABHD12 knockout (–/–) mice following intraperitoneal injection of LPS (10 mg/kg body weight, 4 hours), showing increased average size of activated microglia from the ABHD12 knockout mice in every biological replicate tested. The "n" in parenthesis represents number of activated cerebellar microglial cells quantified per biological replicate. ***p < 0.001 versus (+/+) group by Student's two-tailed unpaired parametric *t*-test;

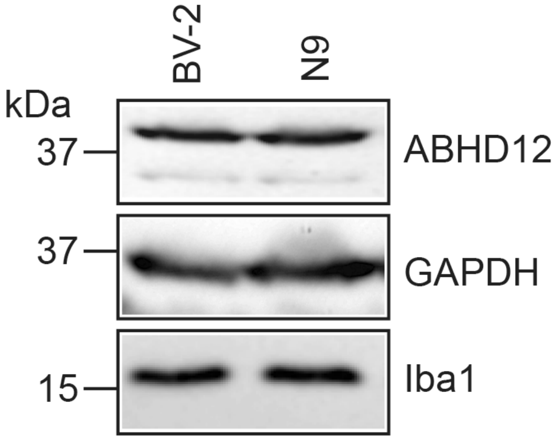


**Figure S2. ABHD12 is expressed in the BV-2 and N9 microglial cell lines.** Western blot analysis showing the expression of ABHD12 in the BV-2 and N9 microglial cells. In this experiment, Iba-1 and GAPDH were used as loading controls. This western blot experiment was done two times (two separate biological replicates) with reproducible results both times.

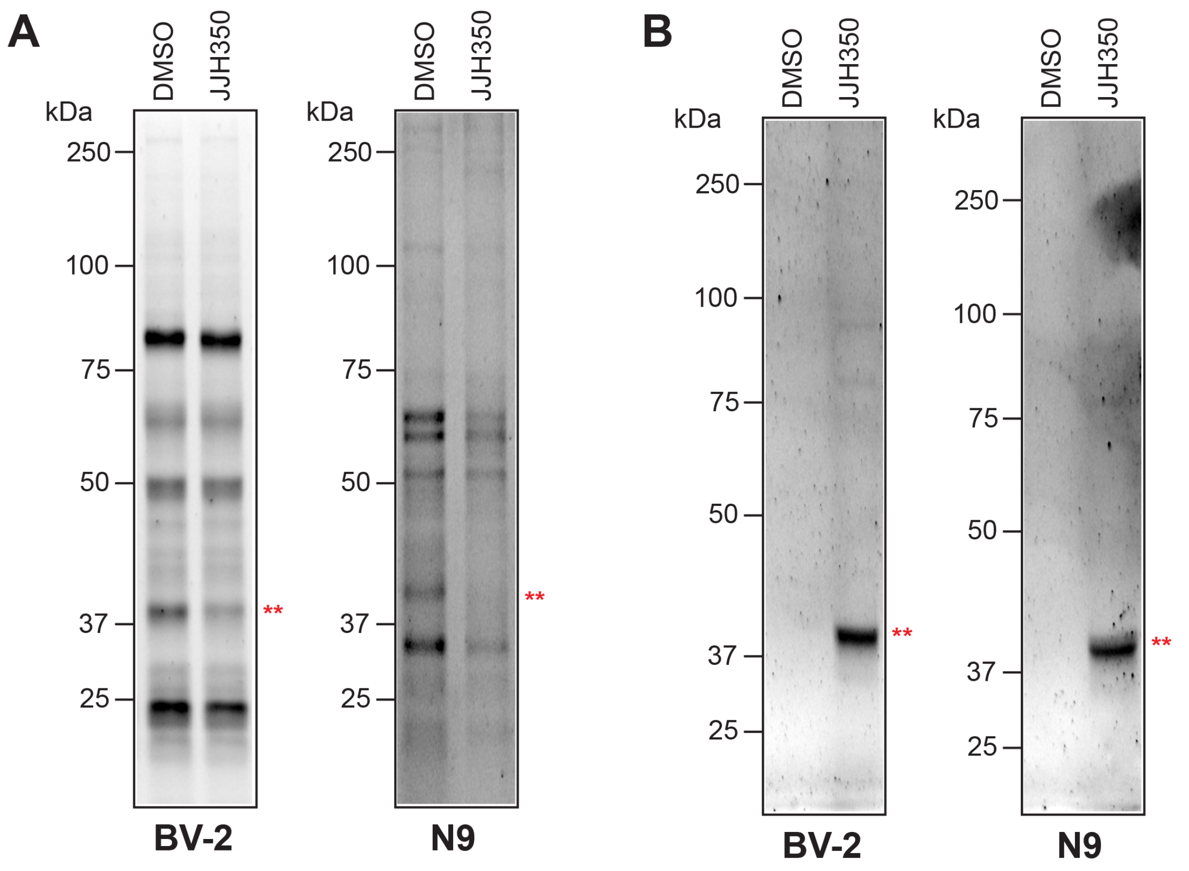


**Figure S3. Competitive gel-based activity based protein profiling (ABPP) assays showing inhibition of ABHD12 by JJH350 in microglial cells**. (**A**) Competitive gel-based ABPP assays on membrane lysates of BV-2 and N9 microglial cells treated with DMSO or JJH350 (10 $\mu$M, 6 hours) using the serine-hydrolase directed FP-rhodamine probe (2 $\mu$M, 45 mins) showing loss of ABHD12 following JJH350 treatment. (**B**) Click reactions on membrane lysates of BV-2 and N9 microglial cells treated with DMSO or JJH350 (10 $\mu$M, 6 hours) showing that JJH350 binds selectively to ABHD12 as per literature reports. Both experiments were done thrice (biological replicates) with reproducible results each time. The red double asterisk on all the ABPP gels correspond to the ABHD12 activity band.

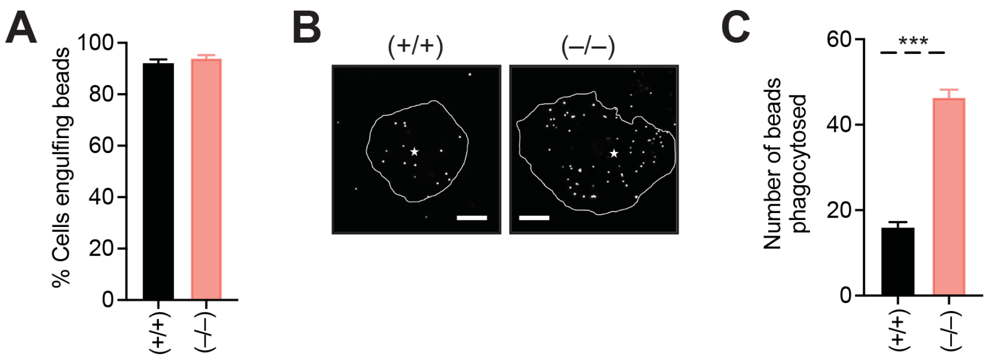


**Figure S4. Loss of ABHD12 results in increased phagocytosis by primary peritoneal macrophages in a fluorescent latex bead uptake assay**. (**A**) Quantification of the total percentage of primary thioglycollate-elicited peritoneal macrophages engulfing fluorescent latex beads following vehicle (DMSO) or JJH350 (10 μM, 6 hours) treatment, showing no significant differences between the two experimental groups. Data (bar plots) is represented as mean ± standard deviation from six independent experiments (biological replicates) per experimental group. (**B**) Representative microscopy images of primary thioglycollate-elicited peritoneal macrophages engulfing fluorescent beads following vehicle (DMSO) or JJH350 (10 μM, 6 hours) treatment, showing increased phagocytosis of fluorescent beads in the JJH350-treated primary thioglycollate-elicited peritoneal macrophages. Scale bar in the microscopy images is 10 μm and (*) represents centroid of the nucleus. (**C**) The number of fluorescent beads phagocytosed by primary thioglycollate-elicited peritoneal macrophages following vehicle (DMSO) or JJH350 (10 μM, 6 hours) treatment, showing a significant increase of number of phagocytosed fluorescent latex beads by the JJH350-treated primary thioglycollate-elicited peritoneal macrophages. Data (bar plots) is represented as mean ± standard deviation from six independent experiments (biological replicates) per experimental group. ***$p < 0.001$ versus vehicle group by Student's two-tailed unpaired parametric *t*-test.
